# A widespread bacterial toxin reveals a deeply divergent DNase family neutralized by diverse immunity proteins

**DOI:** 10.64898/2026.09.22.753552

**Authors:** Audrey R. Long, Gemma Simpson, Julia T. Hespanhol, Kristi Pham, Andrew L. Lovering, Ethel Bayer-Santos

**Author notes:** Corresponding authors (EBS); (ALL).

## Abstract

Bacteria compete for space and resources by delivering toxin proteins into neighboring cells. Many toxins remain uncharacterized because they lack similarity to proteins of known function. Here we identify Rehh as a widespread antibacterial toxin associated with multiple bacterial secretion systems and show that it is used in interbacterial competition. Purified recombinant Rehh functions as a manganese-dependent DNase, consistent with our *in-vivo* findings that Rehh damages target cell DNA, inducing the SOS response and the accumulation of double-strand breaks. The crystal structure of Rehh in complex with a cognate immunity protein reveals a positively charged histidine-rich metal-binding pocket embedded within a protein scaffold that bears no significant similarity to previously characterized nucleases. However, local structural analysis demonstrates that this pocket preserves the catalytic geometry characteristic of the HNH/His-Me-finger superfamily, revealing Rehh to be a deeply divergent HNH nuclease whose ancestry has been obscured through extensive structural remodeling. The cognate immunity protein neutralizes the toxin via an acidic surface that occludes the catalytic pocket, consistent with mimicry of the electrostatic properties of the DNA phosphate backbone. Comparative genomics further shows that Rehh homologs are associated with multiple unrelated immunity families, suggesting that distinct immunity domains repeatedly converge on a common strategy for inhibition. Together, these findings demonstrate how conflict-driven diversification can obscure enzyme ancestry while preserving catalytic function and reveal a possible case of convergent evolution among immunity proteins that independently exploit the same toxin biochemical vulnerability.

**Highlights:**

- Rehh is a widely distributed antibacterial toxin delivered by several secretion systems
- Rehh intoxication causes DNA damage and activates the bacterial SOS response
- Rehh is a manganese-dependent DNase with a deeply divergent HNH active site
- Different families of structurally unrelated immunity proteins neutralize Rehh via acidic DNA-mimicking surfaces

## INTRODUCTION

Bacteria compete for limited space and resources by delivering toxin proteins directly into neighboring cells^1,2^. These antibacterial toxins encompass extensive biochemical diversity^3,4^ but many remain uncharacterized because their sequences and predicted structures lack detectable similarity to proteins of known function^5^. Consequently, a substantial portion of bacterial toxin diversity remains hidden within genomic databases^6,7^. Defining the activities and evolutionary relationships of these proteins is required both to understand full ecological dynamic of interbacterial competition and to reveal how conserved biochemical functions persist during extensive protein diversification.

Protein function can be preserved long after sequence similarity and even global structural similarity have eroded beyond confident recognition by current annotation methods^8^. In such cases, evolutionary relationships may be detectable primarily through the local organization of catalytic residues and their associated secondary structural elements rather than through overall protein architecture^3,9^. The HNH/His-Me-finger nuclease superfamily illustrates this principle^10,11^. Members of this superfamily share a compact ββα-metal catalytic core related to the treble clef motif^12^. A divalent metal ion is typically coordinated by conserved histidine and/or asparagine residues, while a separate histidine promotes nucleophilic attack on the scissile phosphate^11,13,14^. Because catalysis depends primarily on the spatial arrangement of a small number of residues within the HNH core architecture, the surrounding scaffold can undergo extensive remodeling while retaining nuclease function^10,11^. In many cases, large insertions and accessory structural elements accumulate between the canonical β-strands and α-helix of the treble clef motif^10,12^. As a result, deeply divergent HNH nucleases may escape annotation by sequence comparison or whole-structure searches even when their defining catalytic geometry remains intact.

Such divergent proteins are particularly common within polymorphic toxin systems. In these conflict systems, conserved delivery modules are repeatedly combined with rapidly evolving toxin domains that are exchanged among diverse secretion architectures^3,4^. This modular organization enables toxin repertoires to diversify independently of their delivery machinery and facilitates the continual emergence of new toxin variants. Nucleases are among the most abundant and repeatedly recruited enzymatic activities in these systems^3–5,15^. Consequently, polymorphic toxins represent a rich source of protein families in which core catalytic mechanisms may be preserved despite extensive sequence and structural divergence.

Polymorphic toxin systems are typically accompanied by cognate immunity proteins encoded adjacent to toxin genes that protect producing cells from self-intoxication^3,4^. Non-cognate immunity systems, which can also be acquired independently and maintained as orphan defense loci, have been reported to contribute to protection against interbacterial attack^3,13,16–18^. Comparative genomic and experimental studies have revealed complex evolutionary relationships between toxins and their inhibitors. For example, members of the SUKH superfamily are associated with diverse nuclease and nucleic-acid deaminase toxins, demonstrating that a single immunity superfamily can be adapted to recognize multiple classes of toxin partners^15^. Conversely, RNase toxins can recruit structurally distinct immunity proteins, indicating that conservation of toxin architecture does not require co-conservation of the corresponding immunity scaffold^19^. Together, these observations highlight the evolutionary flexibility of toxin-immunity partnerships.

Here, we characterize Rehh, a previously uncharacterized toxin family that is widely distributed across bacterial phyla and associated with multiple protein secretion systems. Rehh (formerly designated STox_30) was identified through a systematic survey of unannotated polymorphic toxin domains across 10,000 *Salmonella* genomes, which revealed nucleases as a major class of previously uncharacterized toxin activities^5^. We sought to determine the biological activity of this family, define its catalytic mechanism, and understand how it is recognized by cognate immunity proteins. Our findings reveal an unrecognized connection between Rehh and the HNH nuclease superfamily and uncover unexpected diversity in the immunity proteins associated with this toxin lineage. More broadly, this work illustrates how bacterial conflict systems can conceal deeply divergent members of established enzyme superfamilies and provides insight into the evolutionary strategies used to maintain toxin neutralization despite extensive diversification.

## RESULTS

### Rehh is a polymorphic toxin encoded across diverse bacterial secretion systems

A systematic survey of unannotated polymorphic toxins previously identified a C-terminal domain, formerly designated STox_30, fused to an N-terminal PAAR domain via RHS repeats^5^. This domain showed no detectable sequence or predicted structural similarity to any previously characterized protein family, indicating that it represents a novel toxin family. We renamed this domain Tox-REHH, reflecting the conserved active-site residues identified in this study (Arg, Glu, His, His; see below). To determine whether Tox-REHH is broadly distributed, we searched for homologs across sequenced bacterial genomes and found that Tox-REHH is encoded across numerous bacterial phyla, spanning both Gram-positive and Gram-negative lineages (Fig. 1A). For detailed mechanistic characterization, we selected a single representative homolog from *Salmonella enterica*, hereafter referred to as Rehh, together with its cognate immunity protein RehhI, which contains an identifiable Imm22 domain^3^.

**Figure 1.**
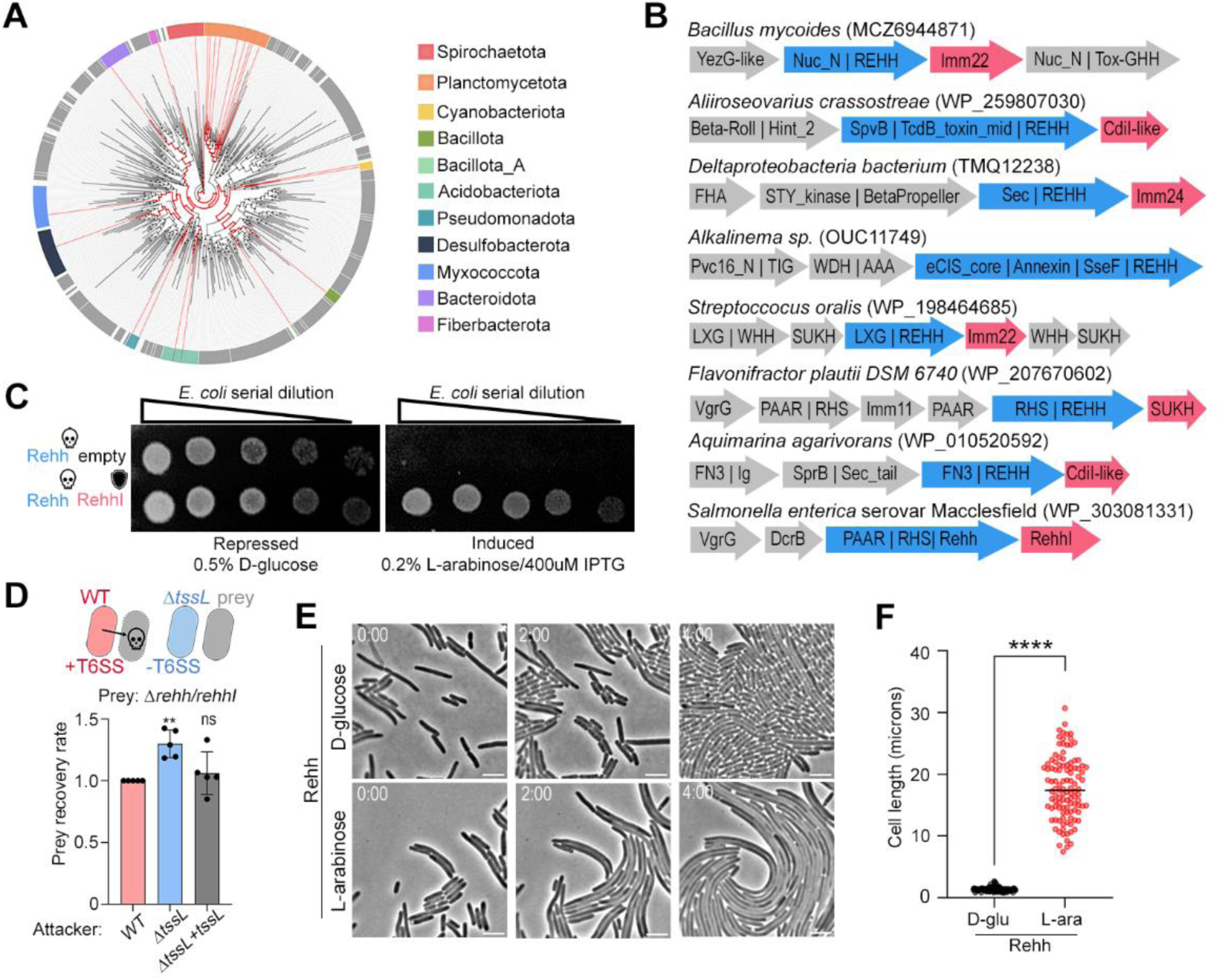
Tox-REHH is a widespread antibacterial toxin deployed by diverse secretion systems. A) Taxonomic distribution of Tox-REHH homologs across bacterial phyla. B) Representative genomic contexts showing association of Tox-REHH homologs with distinct secretion systems. C) Toxicity assay in *E. coli*. Expression of Rehh inhibits growth, whereas co-expression of the cognate immunity protein RehhI restores viability. D) T6SS-dependent interbacterial competition assay. Wild-type *Salmonella* attacker cells outcompete a Δ*rehh-rehhI* prey strain, whereas deletion of *tssL* abolishes antagonism. Complementation with *tssL* restores competitive activity. Data reflects mean ± SD. Statistical significance was assessed by one-way ANOVA followed by Dunnett’s multiple comparison test. **p<0.01. E) Time-lapse microscopy of *E. coli* expressing Rehh under inducing or repressing conditions over 4 h. Rehh expression causes progressive cell elongation. Scale bar, 5 μm. See Movies S1 and S2. F) Quantification of cell length after 4 h. One hundred cells were measured per condition. Bars represent mean ± SD. ****P < 0.0001 (Welch’s t-test).

We next examined the genomic neighborhoods of Rehh homologs to identify the delivery systems and candidate immunity genes with which the toxin is associated (Fig. 1B). In Gram-positive bacteria, Rehh is frequently fused to an LXG or WXG100 domain and encoded alongside components of the type VII secretion system^20^ or fused to a non-catalytic Nuc_N domain within gene clusters encoding other nucleases^15^. In Gram-negative bacteria, Rehh homologs are associated with the type V secretion system^21^ and with the type VI secretion system^22^ via the RHS. A smaller number of homologs are associated with the type IX secretion system^23,24^, a Tc toxin delivery system^25^, and the *Photorhabdus* virulence cassette^26^.

To determine whether Rehh functions as an antibacterial toxin, we expressed the toxin domain in *Escherichia coli* in the presence or absence of its cognate immunity protein, RehhI. Induction of Rehh in the absence of RehhI arrested growth, whereas cells co-expressing RehhI, or cells in which Rehh expression was repressed by glucose, grew normally (Fig. 1C). These results establish Rehh as a potent antibacterial toxin and demonstrate that RehhI is sufficient to neutralize its activity.

Because Rehh is encoded within a T6SS-associated locus in *Salmonella*, we next asked whether it functions as a naturally deployed effector during interbacterial competition. Wild-type attacker cells efficiently outcompeted target bacteria lacking the *rehh-rehhI* locus, whereas deletion of the essential T6SS structural component *tssL* abolished this competitive advantage (Fig. 1D). Complementation of the Δ*tssL* strain restored antagonistic activity, confirming that the observed phenotype depends on a functional T6SS. Together, these results demonstrate that Rehh is delivered during T6SS-mediated interbacterial competition and contributes to the antagonism of neighboring bacterial cells.

We further investigated the cellular consequences of Rehh intoxication using time-lapse microscopy. Cells expressing Rehh progressively elongated over time during induction with arabinose, whereas repressed cells maintained a normal rod-shaped morphology (Fig. 1E; Movie S1; Movie S2). Quantification of cell length confirmed that Rehh-expressing cells were significantly longer than repressed controls (Fig. 1F). Because filamentation is a hallmark of impaired cell division and is commonly associated with activation of the bacterial SOS response^27–29^, this phenotype raised the possibility that Rehh intoxication triggers a DNA-damage response, a hypothesis we test directly below.

Together, these results establish Rehh as an antibacterial toxin actively deployed during interbacterial conflict and show that Tox-REHH more broadly defines a polymorphic toxin family widely distributed across bacterial phyla and repeatedly associated with distinct secretion systems.

### Rehh contains a conserved positively charged metal-binding pocket occluded by an acidic RehhI interface

To gain structural insight into Rehh function, we solved the crystal structure of the Rehh– RehhI complex at 2.5 Å resolution (Fig. 2A). The overall Rehh architecture does not show convincing global similarity to any previously characterized protein in the PFAM database^30^, consistent with its assignment to the “not determined” category in a previous study^5^. However, mapping sequence conservation from an alignment of Rehh homologs onto the structure revealed that the most highly conserved residues cluster within a single spatially discrete pocket (Fig. 2A– B). Five residues within this pocket (R46, H52, E53, H102, and H121) are conserved across Rehh homologs, and individual point mutation of four of these residues abolished toxicity in our heterologous expression assay, indicating that this pocket is required for Rehh activity/toxicity (Fig. 2C). The structure reveals that the three conserved histidine residues (H52, H102, and H121) coordinate a divalent metal ion within this pocket, which carries a strongly positive electrostatic potential (Fig. 2B, left).

**Figure 2.**
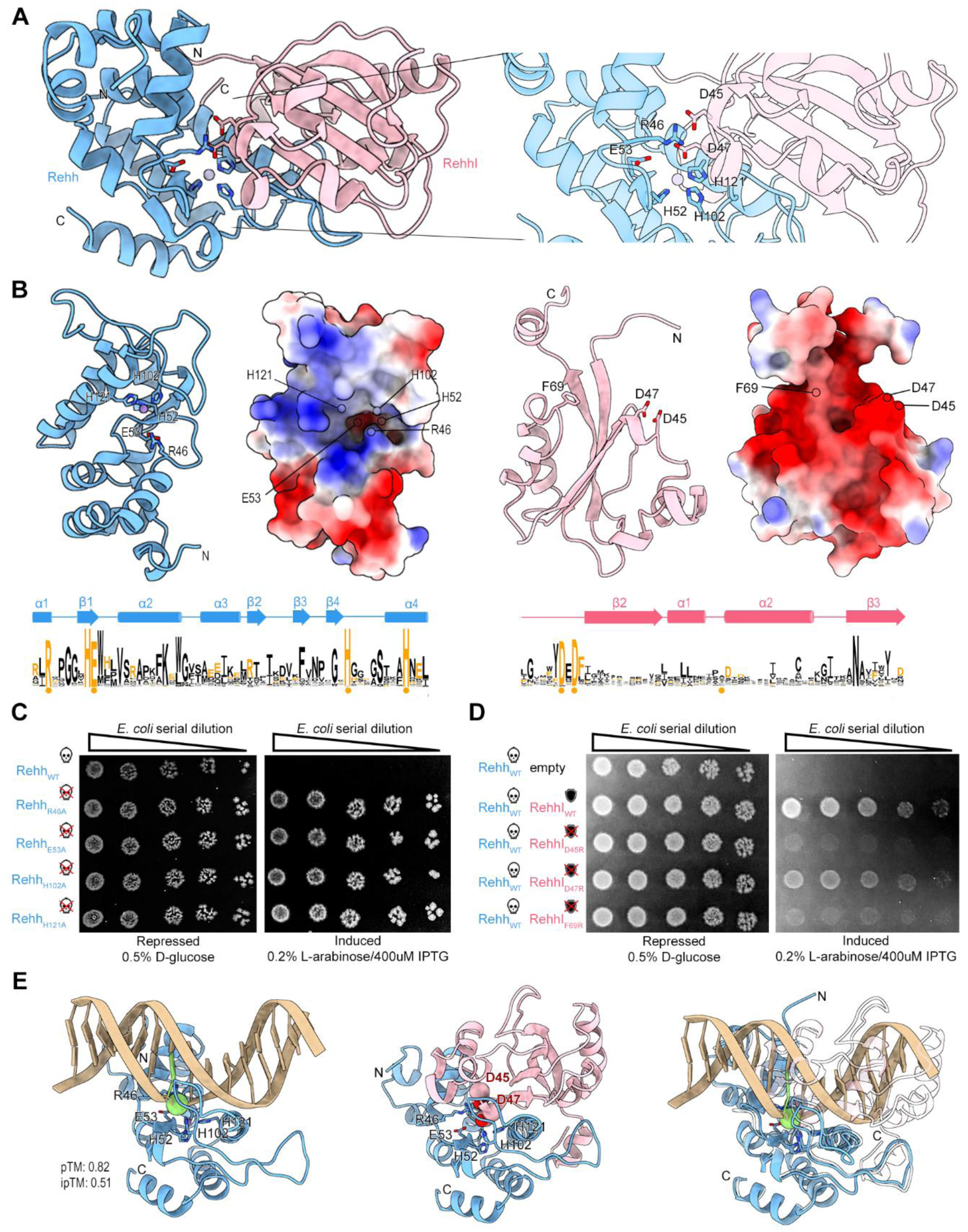
The Rehh-RehhI complex reveals a conserved metal-binding active site inhibited by an acidic immunity interface. A) Crystal structure of the Rehh-RehhI complex (left) and enlarged view of the toxin-immunity interface (right). Conserved Rehh residues (R46, H52, E53, H102, and H121) cluster within a metal-binding pocket and interact with conserved RehhI residues. B) Electrostatic surface representation of Rehh and RehhI. Conserved residues from Tox-REHH and Imm22 family alignments are indicated. C) Toxicity assays of Rehh active-site variants. Mutation of conserved residues R46, E53, H102, or H121 abolishes toxicity. D) Neutralization assays of RehhI interface mutants. Mutations D45R, D47R, and F69R reduce or disrupt protection against Rehh-mediated toxicity. E) Structural model of double-stranded DNA positioned within the Rehh active-site pocket. The DNA phosphate backbone occupies a region similar to that contacted by conserved acidic residues of RehhI. Structures were visualized using ChimeraX^31^.

We next examined how RehhI neutralizes the toxin. In the structure of the complex, a surface feature of RehhI directly overlays this positively charged Rehh pocket (Fig. 2A). This immunity surface is acidic with two conserved residues (D45 and D47), forming electrostatic contacts with the conserved active-site residues of Rehh (Fig. 2A). A hydrophobic contact between RehhI residue F69 and Rehh further stabilizes the complex interface, although this residue is not strictly conserved among RehhI homologs (Fig. 2B, right). Charge-reversal mutations at the immunity interface (RehhI_D45R_ and RehhI_F69R_) most strongly disrupted the ability of RehhI to neutralize Rehh toxicity, whereas RehhI_D47R_ had a smaller effect with residual protection still observed (Fig. 2D). These results indicate that both the electrostatic and hydrophobic components of the RehhI–Rehh interface contribute to inhibition.

The observation that RehhI neutralizes Rehh by presenting an acidic surface to a positively charged metal-coordinating pocket raised the possibility that this surface mimics the electrostatic properties of a nucleic acid substrate. Consistent with this idea, cells intoxicated with Rehh progressively elongate (Fig. 1E), a phenotype often associated with the SOS response to DNA damage^27^. To test whether the geometry of the Rehh pocket is compatible with nucleic acid binding, we co-folded a double-stranded DNA substrate into the predicted active site of the toxin using AlphaFold^32^. DNA fits favorably within the pocket, with its phosphate backbone positioned close to the conserved histidines and metal ion (Fig. 2E). Notably, the two most conserved RehhI residues, D45 and D47, occupy a position equivalent to that of the DNA phosphate backbone in this docking model (green sphere in Fig. 2E).

Together, these structural and mutational data indicate that RehhI neutralizes Rehh by presenting an acidic surface that occludes a positively charged, metal-binding pocket capable of accommodating DNA. We favor a model in which RehhI mimics the electrostatic properties of the DNA phosphate backbone substrate to competitively block this site. We next tested whether Rehh functions as DNase.

### Rehh is a manganese-dependent DNase that activates the SOS response

To test whether Rehh intoxication activates the SOS response, we monitored expression GFP under the control of *recA* promoter^33^ in cells expressing either the Rehh_wt_, the catalytically inactive point mutant Rehh_H121A_, or an empty vector control. GFP fluorescence increased specifically in cells expressing Rehh_wt_ and was not observed in cultures repressed by glucose or in cells expressing Rehh_H121A_ (Fig. 3A). Because the *recA* reporter is activated by the SOS response, these data indicate that Rehh activates this pathway in a manner that depends on an intact toxin active site.

**Figure 3.**
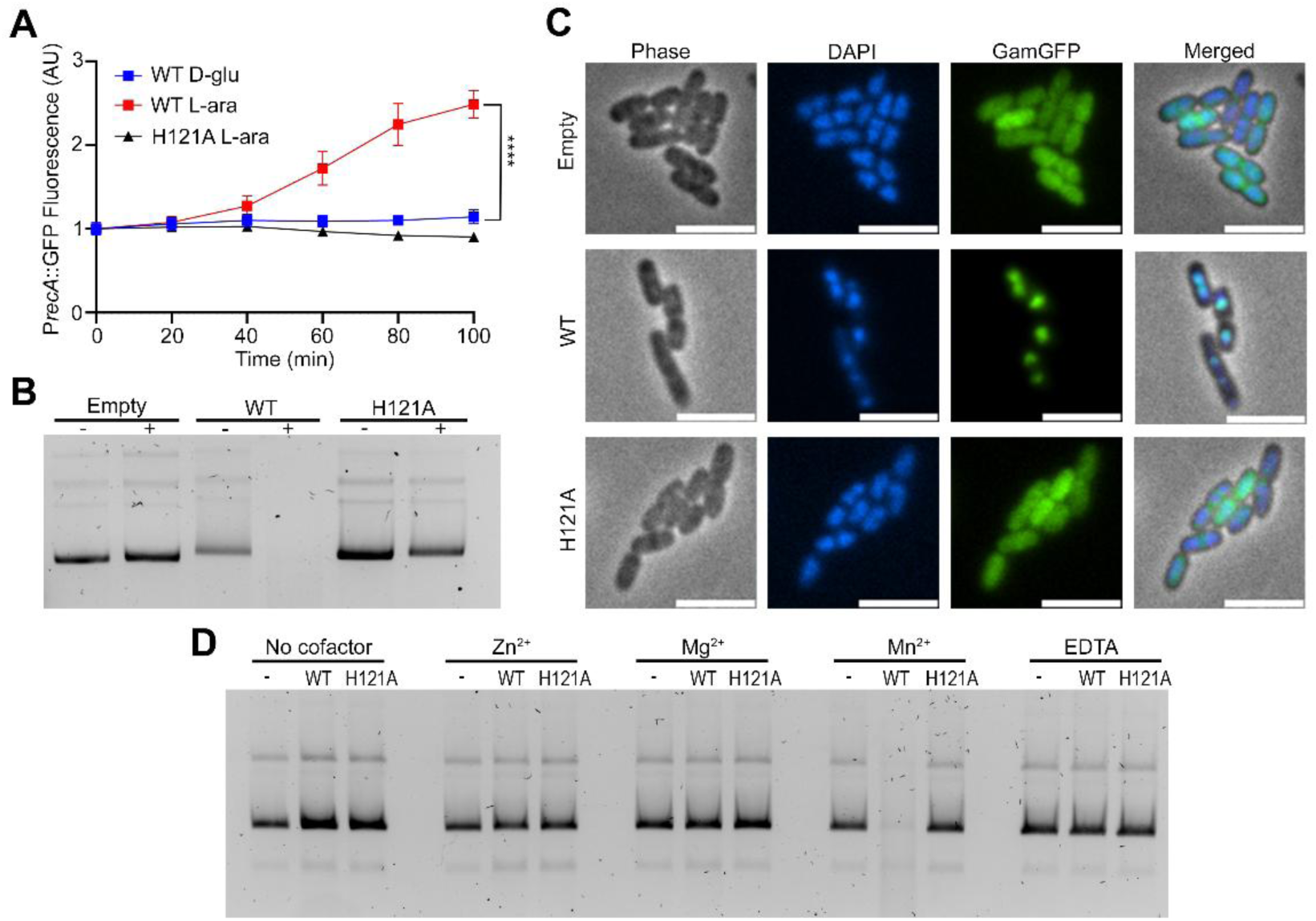
Rehh induces DNA damage and functions as a manganese-dependent DNase. A) SOS-response reporter assay using a P*recA*-GFP transcriptional fusion. Expression of Rehh_wt_, but not Rehh_H121A_ or the empty-vector control, activates GFP expression. Points represent mean ± SD from three biological replicates. ****P < 0.0001 (Welch’s t-test). B) Agarose gel analysis of plasmid DNA recovered from cells expressing Rehh_wt_, Rehh_H121A_, or an empty vector. Expression of Rehh_wt_ reduces recoverable plasmid DNA. C) Fluorescence microscopy of GamGFP, a reporter for double-strand DNA breaks, in cells expressing Rehh_wt_, Rehh_H121A_, or the empty vector control. DAPI staining identifies nucleoids. D) *In vitro* DNase assay using purified recombinant protein. Rehh_wt_ degrades plasmid DNA in the presence of Mn²⁺, whereas Rehh_H121A_ is inactive.

Because the SOS response is triggered by DNA damage^34^, we asked whether Rehh intoxication is accompanied by degradation of cellular DNA. We extracted plasmid from cells expressing an empty vector, Rehh_wt_ or Rehh_H121A_ after one hour of induction. Recoverable plasmid DNA was depleted specifically in cells expressing Rehh_wt_, whereas plasmid yield was unaffected in the repressed condition or in cells expressing the catalytically inactive mutant (Fig. 3B). Total RNA extracted from the same cells did not show any sign of degradation, suggesting that Rehh activity is specific to DNA rather than a general nuclease (Fig. S1).

To visualize DNA damage directly in cells, we used fluorescence microscopy with cells expressing a fusion of GFP to Gam protein from the bacteriophage Mu (GamGFP), which binds the ends of double-stranded DNA breaks and forms discrete foci at sites of cleavage^35^. Cells expressing Rehh_wt_ developed distinct GamGFP foci that were absent in cells expressing an empty vector or Rehh_H121A_ (Fig. 3C). GamGFP foci consistently overlapped with regions stained with DAPI, indicating that the signal detected by GamGFP occurs within regions containing nucleoid DNA. Together, these results demonstrate that Rehh intoxication damages cellular DNA, including the accumulation of double-strand breaks, in a manner that depends on an intact toxin active site.

To determine whether Rehh acts directly on DNA, we purified recombinant Rehh and analyzed whether it is sufficient to degrade DNA *in vitro*. Because Rehh is co-expressed with RehhI to avoid toxicity during expression, we first purified the Rehh–RehhI complex, then used a denaturation–refolding strategy to separate Rehh from the immunity protein (Fig. S2). Because the Rehh structure revealed three conserved histidines coordinating a divalent metal ion within the active-site pocket (Fig. 2A), we also tested whether Rehh activity depends on a specific metal cofactor. Purified Rehh_wt_ was dialyzed to remove any traces of metal and incubated with plasmid DNA in the presence of Mg^2+^, Mn^2+^, Zn^2+^or EDTA. Under the conditions tested, detectable Rehh DNase activity required Mn^2+^; no degradation was observed with Mg^2+^, Zn^2+^, or EDTA (Fig. 3D), although some degradation was observed with Mg^2+^ during longer incubation periods. Together, these results establish that Rehh functions as a manganese-dependent DNase.

We next sought to determine any Rehh evolutionary relationship to previously characterized nucleases. The conserved histidine-rich active site and coordinated divalent metal ion suggested similarity to HNH nucleases, a large superfamily of metal-dependent DNases that includes numerous polymorphic toxins^3,10^. Although HNH nucleases are highly diverse in sequence and overall structure, they share a conserved catalytic architecture derived from a treble clef motif. This motif is defined by three structural elements: a Zn-binding knuckle, a β-hairpin, and an α-helix^12^. Although the Zn-binding knuckle is not always retained, alternative metal-binding residues are typically acquired when this element is lost^12,36^.

Rehh contains these defining structural elements (Zn-knuckle, M51-S79; β-hairpin, W84-P101; α-helix R115-Q130) despite lacking detectable sequence similarity or obvious global structural similarity to known HNH nucleases. The conserved catalytic residues identified by mutagenesis are positioned mostly within a β-hairpin–α-helix arrangement that coordinates a divalent metal ion in a geometry closely resembling established HNH active sites (Fig. S3). In canonical HNH nucleases, a conserved catalytic Arg residue is typically contained within this ββα-element. However, the conserved R46 residue of Rehh lies just upstream of the ββα rather than within it. This difference may have resulted from the adjacent insertions unique to this subfamily. Like many toxin-associated HNH nucleases^3^, Rehh also contains substantial insertions within and surrounding the treble clef motif, which likely contribute to the divergence that prevented its recognition. Together, these data support experimentally-led classification of the Tox-REHH domain as a deeply divergent member of the HNH superfamily.

### Rehh homologs are neutralized by structurally diverse immunity domains that share a negative electrostatic surface

Having established that RehhI neutralizes Rehh by occluding its positively charged active-site pocket, we next asked whether this mode of neutralization is conserved across the broader Tox-REHH family. Examination of the genomic neighborhoods of Rehh homologs revealed several cases in which the toxin was associated with candidate immunity genes encoding domains distinct from the Imm22 of RehhI (Fig. 1B). Associations between a toxin family and multiple immunity-domain families have been reported previously for a limited number of polymorphic toxins but this observation remains with little characterization to date^3^. We therefore used AlphaFold to predict the structures of these toxin-immunity complexes and assess whether the different immunity proteins engage the toxin through a common mechanism (Fig. 4).

**Figure 4.**
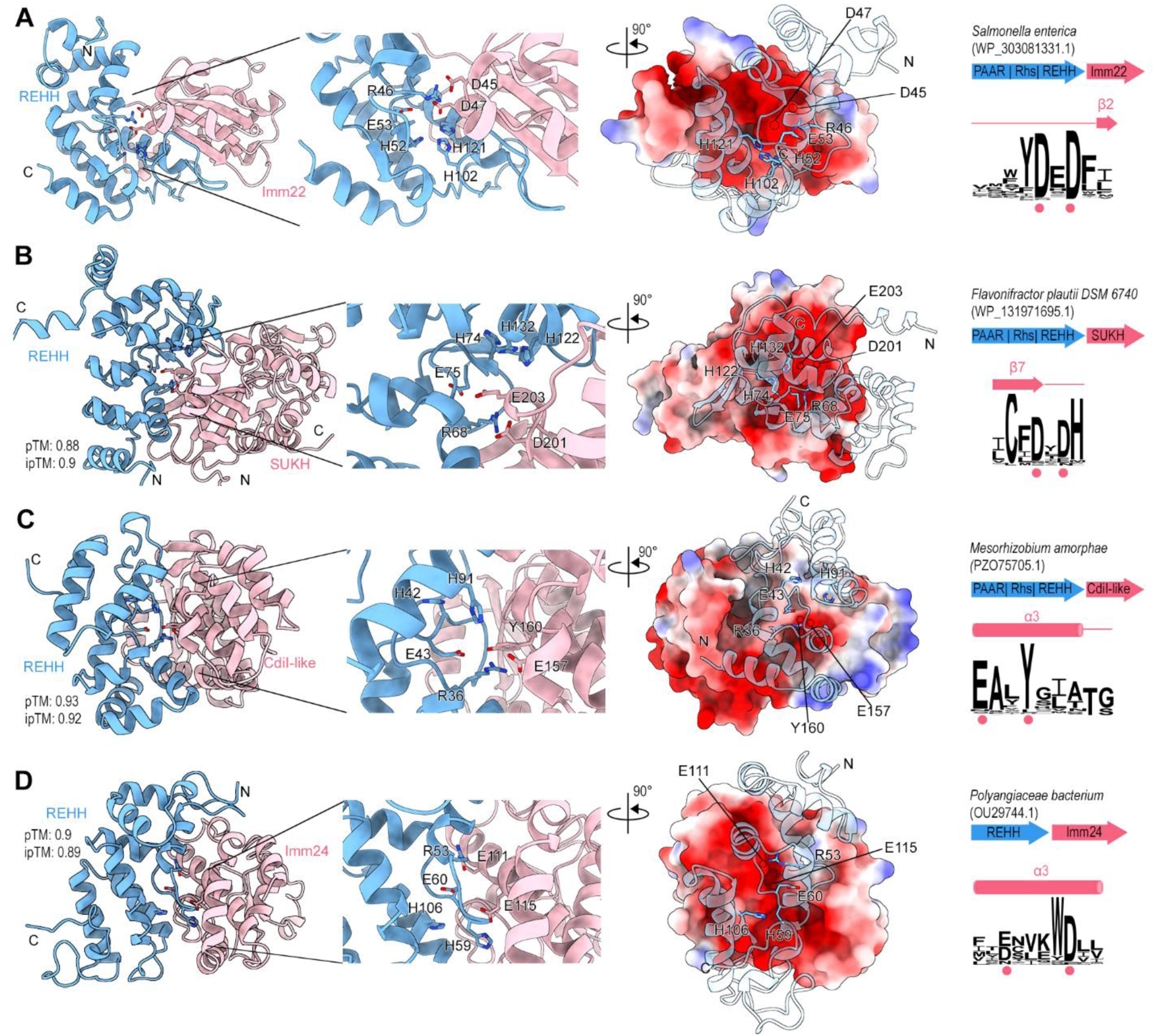
Distinct immunity families neutralize Tox-REHH toxins through conserved acidic interfaces. A) Experimentally determined Rehh-RehhI complex showing inhibition of the conserved toxin active site by the Imm22 immunity protein. B) AF3 model of a Tox-REHH homolog bound to a SUKH-domain immunity protein. C) AF3 model of a Tox-REHH homolog bound to a CdiI-like immunity protein. D) AF3 model of a Tox-REHH homolog bound to an Imm24 immunity protein. Electrostatic surface representations highlight conserved acidic residues positioned toward the toxin active site in each complex. Predicted structures were visualized using ChimeraX^31^. The genomic organization of the toxin-immunity locus and sequence logo of the associated immunity family are shown on the right.

In addition to Imm22, we identified several other immunity families associated with Rehh homologs, including SUKH, CdiI-like, and Imm24^3,15^. Although these domains are structurally unrelated to one another, all are predicted to bind the same conserved surface of the toxin occupied by RehhI (Fig. 4A-D and Fig. S4). In each case, conserved acidic residues are positioned toward the positively charged catalytic pocket of the toxin, despite being presented on different secondary structural elements. These conserved electronegative elements of the different immunity proteins are predicted to occupy the same active-site position as the DNA backbone phosphate, as we observed with Rehh-RehhI (Fig. S5). However, there are some differences at the interface of the predicted toxin-immunity complexes. In the cases of the Rehh homologs that pair with CdiI-like and Imm24 immunity proteins, the toxin active site undergoes a conformational change after binding of the immunity protein when compared to the predicted structure of the toxin alone (Fig. S6).

Together, these analyses indicate that a conserved positively charged toxin surface is repeatedly targeted by multiple unrelated immunity families. The recurrence of acidic interaction surfaces directed toward the same catalytic pocket suggests that distinct immunity lineages have converged on a common strategy for neutralizing Tox-REHH toxins. Notably, this conserved electrostatic arrangement mirrors the DNA-mimicry mechanism proposed for RehhI (Fig. 2E), raising the possibility that structurally unrelated immunity proteins independently exploit the nucleic acid-binding properties of the toxin to achieve inhibition.

## DISCUSSION

Bacterial conflict systems continue to reveal unexpected diversity in even the best-characterized classes of proteins. Here, we show that Rehh defines a previously unrecognized branch of the HNH superfamily and provide evidence that structurally unrelated immunity proteins can repeatedly converge on a common strategy for neutralizing this toxin family. These findings highlight how conflict-driven evolution can simultaneously obscure the ancestry of toxin domains and promote diversification of their immunity partners.

The difficulty in recognizing Rehh as an HNH nuclease likely reflects a general feature of the HNH superfamily itself. Catalysis is mediated by a compact treble clef core motif whose function depends on the spatial arrangement of only a small number of residues, while the surrounding scaffold can tolerate extensive structural remodeling^10–12^. As a result, sequence conservation and even global structural similarity may erode beyond recognition while catalytic geometry remains intact. Rehh appears to represent an extreme example of this phenomenon, with extensive elaboration of the ancestral architecture obscuring its relationship to previously characterized HNH nucleases. More broadly, our findings highlight a limitation of current annotation approaches, which frequently rely on conservation across entire domains and may therefore overlook proteins whose evolutionary signal is restricted to a small catalytic core.

Polymorphic toxin systems are likely to be especially rich reservoirs of such hidden diversity. These systems evolve under intense selective pressure imposed by interbacterial competition, driving continual diversification of toxin repertoires while preserving biochemical function^3,4^. Diversification is favored because toxin efficacy depends on overcoming the immunity proteins and resistance mechanisms present in neighboring competitors. As immunity proteins diversify and spread through bacterial populations, toxin variants that avoid immunity recognition while retaining catalytic activity can escape pre-existing protection and provide a transient competitive advantage to the producer cell^4,18,36–38^. Reciprocally, selection favors the emergence of new immunity proteins capable of neutralizing these toxin variants, generating a continual cycle of toxin and immunity diversification.

Notably, nucleases represent one of the most abundant and repeatedly recruited toxin classes identified in polymorphic toxin systems^3,5,15^. One possible explanation is that nucleic acid cleavage imposes relatively few constraints on overall protein architecture. Unlike toxins that recognize highly specific macromolecular targets (*e.g.* peptidoglycan), nucleases primarily interact with the chemically conserved phosphate backbone of DNA or RNA. This may allow catalytic activity to persist across a broader range of structural scaffolds while tolerating extensive sequence and structural diversification. The deeply divergent architecture of Rehh is consistent with this possibility and suggests that additional unrecognized branches of nuclease superfamilies may remain hidden within polymorphic toxin repertoires.

Structural and computational analyses suggest that RehhI inhibits the toxin by occupying the DNA-binding region of the active site and presenting a negatively charged surface that resembles the electrostatic properties of the DNA phosphate backbone. This strategy is consistent with a broader theme observed across biological conflict systems, where proteins inhibit nucleic-acid-binding enzymes by mimicking features of their natural substrates. One such example is the colicin E9 immunity protein Im9, which binds adjacent to the HNH nuclease active site and blocks DNA binding^39^. Similar principles are also exploited by bacteriophages. The uracil-DNA glycosylase inhibitor Ugi, which presents an extended acidic surface that mimics the shape and charge distribution of duplex DNA, thus occludes the DNA-binding cleft of its target enzyme^40^. The T7 anti-restriction protein Ocr forms a highly negatively charged dimer that structurally resembles bent B-form DNA and protects phage genomes by competitively binding host restriction enzymes^41^. Our results extend this concept to a previously uncharacterized toxin family and suggest that DNA mimicry may represent a broadly accessible solution for neutralizing DNA-targeting enzymes.

The recurrent association of Rehh homologs with multiple unrelated immunity families places this mechanism in an evolutionary context. Comparative genomic studies have shown that individual toxin families can associate with distinct immunity proteins^3^. Previous work on the SUKH superfamily demonstrated that a single immunity fold can be adapted to neutralize multiple toxin classes^15^, while other studies reported closely related RNase toxin families paired with structurally distinct immunity proteins^19^. However, these observations have largely remained descriptive and have lacked a unifying mechanistic explanation. Our data suggest that at least some of these relationships may arise because the primary selective requirement is not conservation of a particular immunity scaffold but neutralization of a conserved biochemical vulnerability. In the case of Rehh, unrelated immunity domains are predicted to converge on the same positively charged toxin surface and to exploit the same electrostatic solution despite adopting different structural architectures. The repeated emergence of acidic toxin-binding interfaces suggests that DNA mimicry may provide a predictable evolutionary route through which new immunity proteins can arise against nuclease toxins.

This model differs from cross-neutralization by non-cognate or orphan immunity systems. Orphan immunity genes can protect against toxins produced by neighboring competitors despite being encoded outside the original toxin-immunity locus^16,17,38^. In most characterized examples, however, protection is mediated by homologs of the same immunity family that have been horizontally acquired and deployed in a different genomic context^38^. By contrast, the immunity proteins associated with Tox-REHH belong to several unrelated immunity families. Rather than diversification within a single immunity lineage, our results suggest repeated recruitment of entirely different protein scaffolds toward a common inhibitory strategy. This distinction may help explain why some toxin families maintain relatively conserved catalytic architectures while simultaneously accumulating remarkably diverse repertoires of associated immunity proteins.

If positively charged nucleic-acid-binding surfaces represent recurrent evolutionary vulnerabilities, then selection may repeatedly favor inhibitors that exploit the same electrostatic principles regardless of their structural origin. Under this model, immunity proteins would not need to evolve toward a specific fold, but rather toward a shared biophysical solution: a negatively charged surface capable of competing with nucleic acid binding. Such a mechanism could help explain both the extraordinary diversity of nuclease toxins and the remarkable plasticity of their immunity partners. More broadly, Rehh illustrates how experimental characterization of unannotated toxin families can uncover hidden branches of ancient enzyme superfamilies while revealing general principles that govern the long-term evolution of toxin-immunity interactions.

## METHODS

### Bacterial strains and growth conditions

Bacterial strains and plasmids used in this study are listed in STAR Methods. Unless otherwise indicated, *E. coli* and *S. enterica* strains were grown in lysogeny broth (LB) at 37°C with aeration. AB defined media was used (0.2% (NH_4_)_2_SO_4_, 0.6% Na_2_HPO_4_, 0.3% KH_2_PO_4_, 0.3% NaCl, 0.1 mM CaCl_2_, 1 mM MgCl_2_, 3 μM FeCl_3_) supplemented with 0.2% sucrose, 0.2% casamino acids, 10 μg/mL thiamine and 25 μg/mL uracil. Antibiotics were used at the following concentrations when required: kanamycin (50 μg/mL), streptomycin (50 μg/mL), chloramphenicol (12.5 μg/mL), tetracycline (10 μg/mL), and ampicillin (100 μg/mL). Expression from the arabinose-inducible P_BAD_ promoter was induced with 0.2% L-arabinose and repressed with 0.2-0.5% D-glucose as indicated. Expression from IPTG-inducible promoters was induced with IPTG concentrations described for individual experiments.

### Bioinformatic analyses

The amino acid sequences of Rehh and RehhI from *S. enterica* serovar Sekondi were used as queries for iterative jackHMMER^42^ searches against the UniProt^43^ and RefSeq^44^ databases. Searches were performed until convergence and retrieval of additional homologs ceased. Homologous sequences were aligned using MAFFT within Jalview^45,46^. Sequence conservation was visualized using WebLogo^47^. Taxonomic distribution of Rehh homologs was visualized using AnnoTree^48^ based on species identified from the homolog collection.

To identify genomic contexts associated with Rehh homologs, curated multiple sequence alignments were converted to profile hidden Markov models and queried against the NCBI nonredundant protein database^49^ using HMMER^50^. Homologs were retrieved using an E-value cutoff of 1 × 10^−2^^5^. Gene neighborhoods were collected and visualized using the ROTIFER package (https://github.com/leepusp/rotifer) with default parameters.

### Cloning, mutagenesis and strain construction

The *rehh* toxin gene and the cognate immunity gene *rehhI* (FD01845383) were synthesized commercially (Twist Bioscience) and cloned into pBRA^51^ and pEXT22^52^ expression vectors, respectively. Site-directed mutants were generated by overlap-extension PCR^53^ and verified by DNA sequencing. Chromosomal deletions in *S. enterica* serovar Sekondi were constructed using λ-Red recombineering^54^. For complementation experiments, *tssL* was cloned into pFPV25.1^55^ under constitutive expression. Primers used in this study are listed in Table S4.

### *E. coli* toxicity assays

Toxicity assays were performed as described previously with minor modifications^56^. Overnight cultures of *E. coli* DH5α carrying pBRA-Rehh variants and, when indicated, pEXT22-RehhI constructs were grown under repressing conditions in LB supplemented with 0.5% D-glucose. Cultures were adjusted to OD600 = 1.0, serially diluted four-fold, and spotted onto LB agar containing either D-glucose or L-arabinose and IPTG to repress or induce toxin and immunity expression, respectively. Images were acquired after overnight incubation at 37°C.

### Interbacterial competition assay

Interbacterial competition assays were performed using *S. enterica* serovar Sekondi as a model T6SS delivery system. Wild-type or Δ*tssL* strains served as attackers, whereas a Δ*rehh-rehhI*::Cm^R^ strain served as prey. Complementation was achieved by expression of *tssL* from pFPV25.1. Overnight cultures were subcultured 1:100 in fresh LB and grown to mid-log phase. Attacker and prey strains were normalized to OD600 values of 4 and 1, respectively, and mixed at equal volumes, resulting in a final attacker-to-prey ratio of 4:1, and spotted onto 0.22-µm nitrocellulose membranes (1×1cm) placed on LB agar supplemented with 0.1% bile salts. Following incubation for 4 h at 37°C, bacteria were recovered, serially diluted, and plated on selective media. Competitive indices were calculated from output-to-input colony-forming unit ratios and normalized to wild-type values.

### Time-lapse microscopy

Time-lapse microscopy was performed using LB agarose pads prepared as described previously^57^. *E. coli* MG1655 carrying pBRA-Rehh was grown overnight under repressing conditions and subcultured to mid-log phase. Cells were normalized to OD600 = 1.0 and deposited onto agarose pads containing either D-glucose or L-arabinose. Images were acquired every 10 min for 4 h using a Nikon microscope equipped with a Prime 95B sCMOS camera and a Plan Fluor 100× Ph3 oil objective. Images were analyzed using Fiji^58^.

### SOS response assay

SOS induction was monitored using *E. coli* MG1655 carrying pSC101-P*recA*::GFP^33^ together with plasmids expressing Rehh_wt_, Rehh_H121A_, or vector controls. Cultures were grown under repressing conditions to mid-log phase, normalized to OD600 = 1.0, and transferred to supplemented AB medium. Toxin expression was induced or repressed by addition of L-arabinose or D-glucose, respectively. Optical density and GFP fluorescence were monitored using a Synergy H1 microplate reader, and fluorescence values were normalized to culture optical density.

### Plasmid degradation assay

To evaluate DNA degradation during intoxication, *E. coli* DH5α carrying empty vector, Rehh_wt_, or Rehh_H121A_ expression constructs were grown to mid-log phase and induced or repressed for 1 h. Plasmid DNA was purified using the GeneJET Plasmid Miniprep Kit and analyzed by electrophoresis on 1% agarose gels containing SYBR Safe. DNA was visualized using a ChemiDoc imaging system.

### Visualization of double-stranded DNA breaks

DNA damage was visualized using the GamGFP DNA break reporter strain SMR1435^35^. Cultures carrying the indicated expression plasmids were grown under repressing conditions to mid-log phase. GamGFP expression was induced with tetracycline for 2 h prior to toxin induction with arabinose for 15 min. Cells were washed with saline buffer, fixed with 4% paraformaldehyde, stained with DAPI, mounted on agarose pads, and imaged using a Nikon microscope equipped with a Prime 95B sCMOS camera and a Plan Fluor 100× Ph3 oil objective. Images were analyzed with Fiji.

### Protein purification and DNase assays

To purify catalytically active toxin, Rehh_wt_ or Rehh_H121A_ was co-expressed with the cognate immunity protein RehhI using pRSF-Duet constructs in *E. coli* BL21(DE3). Rehh variants contained an N-terminal 6×His tag and a C-terminal Strep tag, whereas RehhI was expressed from the second multiple cloning site of the vector with no tag. Overnight cultures were used to inoculate LB medium supplemented with appropriate antibiotics and grown at 37°C to an OD600 ∼0.6. Protein expression was induced with 100 μM IPTG for 4 h. Cells were harvested by centrifugation and lysed by sonication in the presence of lysozyme. The Rehh-RehhI complex was initially purified by affinity chromatography using a 1-mL StrepTrap XT column (Cytiva). The column was equilibrated and washed with buffer containing 50 mM Tris-HCl (pH 8.0), 350 mM NaCl, and 5% glycerol, and bound proteins were eluted with the same buffer supplemented with 50 mM biotin. To obtain toxin free from the cognate immunity protein, pooled fractions containing the Rehh-RehhI complex were applied to a 1-mL HisTrap HP column (Cytiva). The column was washed with buffer containing 50 mM Tris-HCl (pH 8.0), 350 mM NaCl, 5 mM imidazole, and 5% glycerol. Rehh and RehhI were then dissociated by applying a denaturation gradient using buffer supplemented with 8 M guanidine hydrochloride. Following complete denaturation, the toxin was refolded directly on-column by reversing the guanidine gradient. Refolded Rehh was subsequently eluted using an imidazole gradient up to 500 mM imidazole.

Because structural analysis suggested that Rehh coordinates a divalent metal ion through conserved histidine residues, purified toxin preparations were treated to remove copurified metals prior to enzymatic assays. Eluted proteins were buffer exchanged into metal-chelation buffer containing 50 mM Tris-HCl (pH 8.0), 200 mM NaCl, 100 mM EDTA, and 5% glycerol, followed by exchange into storage buffer containing 50 mM Tris-HCl (pH 8.0), 200 mM NaCl, and 5% glycerol. Protein purity was verified by SDS-PAGE.

For DNase assays, purified plasmid DNA was incubated with 50 μM Rehh_wt_ or Rehh_H121A_ in the presence of 1 mM MnCl_2_, MgCl_2_, ZnCl_2_, or EDTA. DNase reactions were performed in storage buffer (50 mM Tris-HCl pH 8.0, 200 mM NaCl, 5% glycerol) and incubated at 37°C for 1 h. Reactions were subsequently analyzed by electrophoresis on 1% agarose gels containing SYBR Safe. DNA was visualized using a Bio-Rad ChemiDoc imaging system.

### Protein crystallization and structure determination

The Rehh-RehhI complex used for crystallographic studies was expressed in *E. coli* Lemo21(DE3). Cells were grown in LB medium supplemented with 100 μM L-rhamnose and antibiotics at 37°C until reaching an OD600 of ∼0.5. Protein expression was induced with 0.5 mM IPTG and cultures were incubated overnight at 25°C with shaking. Cells were harvested by centrifugation and resuspended in buffer A (300 mM NaCl, 50 mM HEPES, 20 mM imidazole, pH 7.5). Following sonication, cell debris was removed by centrifugation at 40,000 *g* for 1 h at 5°C. The clarified lysate was loaded onto a 5-mL HisTrap HP column (Cytiva) pre-equilibrated in buffer A. Bound proteins were eluted using a linear imidazole gradient to buffer B (300 mM NaCl, 50 mM HEPES, 500 mM imidazole, pH 7.5). Fractions containing the Rehh-RehhI complex were concentrated and further purified by size exclusion chromatography on a HiLoad 26/600 Superdex 75 pg column equilibrated in gel filtration buffer (150 mM NaCl, 20 mM HEPES, pH 7.5). Purified protein was concentrated to approximately 16 mg/mL for crystallization trials. Initial crystallization screening was performed by sitting-drop vapor diffusion in 96-well plates using a Mosquito liquid-handling robot (SPT Labtech). Protein and reservoir solutions were mixed at a 1:1 ratio to a final drop volume of 800 nL. Commercial screening kits included Morpheus, MemGold, MemGold2, and JCSG-plus (Molecular Dimensions). Crystals suitable for diffraction grew at 25°C in condition G11 of the MemGold screen containing 0.15 M potassium citrate tribasic monohydrate, 0.05 M lithium citrate tribasic tetrahydrate, 0.1 M sodium phosphate monobasic monohydrate, and 22% (w/v) PEG 6000. Crystals were harvested directly from crystallization drops and flash-cooled in liquid nitrogen.

X-ray diffraction data were collected at beamline ID30B of the European Synchrotron Radiation Facility (ESRF, Grenoble, France). Diffraction images were indexed, integrated, and scaled using XDS through the genades_parallelproc pipeline and further processed using Phenix^59^ and the CCP4 software suite^60^. The structure was solved by molecular replacement using an AlphaFold^32^-predicted Rehh-RehhI complex as the search model. Iterative cycles of manual model building were performed in Coot^61^ followed by crystallographic refinement with REFMAC5^62^. The final structure was refined to 2.4 Å resolution with *R_work_* and *R_free_* values of 0.19 and 0.26, respectively. Complete data collection and refinement statistics are provided in Table S3.

### Statistical analysis

Statistical analyses were performed using GraphPad Prism. Statistical tests used for individual experiments are indicated in the corresponding figure legends. Data are presented as mean ± standard deviation unless otherwise stated.

## ACKNOWLEDGMENTS

This research was supported by startup funds from UT Austin College of Natural Sciences at The University of Texas at Austin to E.B.-S. A.L. and G.S. are supported by Leverhulme Trust Award RPG-2024-336. The authors acknowledge the Texas Advanced Computing Center (TACC) at The University of Texas at Austin for providing computational resources that have contributed to the research reported in this study (http://www.tacc.utexas.edu). The microscopy was performed at the Microscopy & Flow Cytometry Facility at UT Austin (RRID:SCR_021756). We thank the European Synchrotron Radiation Facility (ESRF, Grenoble, France) for provision of synchrotron radiation facilities and access to beamline ID30B. Microsoft Copilot was used to assist with manuscript writing.

## Data and code availability

The crystal structure has been deposited in the Protein Data Bank under accession XXXX. All other data supporting the findings of this study are available from the corresponding author upon request.

