## Supplementary figures for "A widespread bacterial toxin reveals a deeply divergent DNase family neutralized by diverse immunity proteins"

**Supplementary Material**

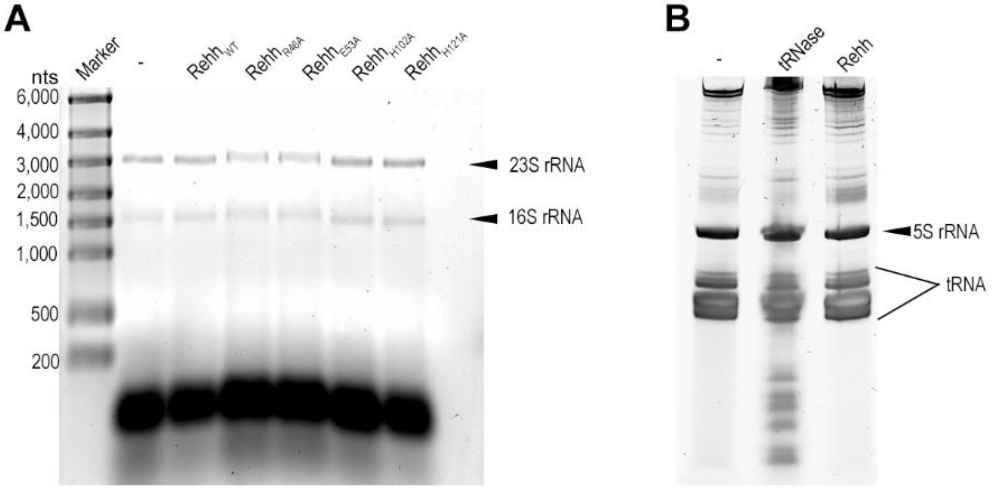


**Figure S1. Rehh has no apparent RNase activity.** A) Total RNA isolated from cells expressing wild-type Rehh, catalytic mutants, or an empty-vector control and analyzed by agarose gel electrophoresis. No degradation of rRNA species was observed. B) Denaturing urea-PAGE analysis of RNA isolated from cells expressing wild-type Rehh. RNA profiles resemble those of the empty-vector control and differ from those produced by a previously characterized tRNase toxin. Related to Figure 3.

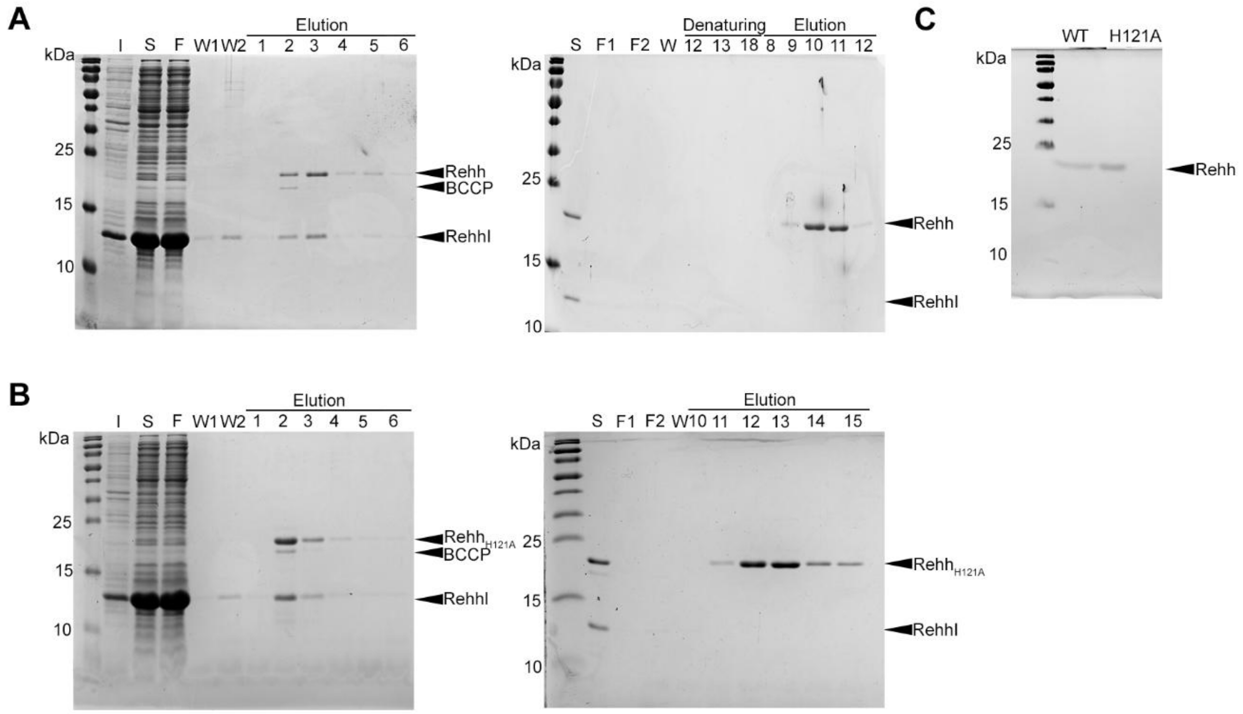


**Figure S2. Purification strategy for recombinant Rehh.** A) Purification of wild-type Rehh. The Rehh-RehhI complex was first isolated by Strep affinity chromatography through the C-terminal Strep tag present on Rehh. Following capture of the toxin-immunity complex, a second purification step using $Ni^{2+}$ affinity chromatography was performed under denaturing conditions to dissociate Rehh from RehhI and isolate the His-tagged toxin. Representative SDS-PAGE analyses of each purification step are shown. B) Purification of $Rehh_{H121A}$ using the same two-step strategy consisting of initial Strep affinity purification of the toxin-immunity complex followed by denaturing $Ni^{2+}$ affinity purification to separate the toxin from RehhI. C) Purified proteins used in biochemical assays after removal of co-purifying metal ions by EDTA treatment and buffer exchange. I, insoluble fraction; S, soluble fraction; F, flow-through; W, wash; E, elution. Related to Figure 3.

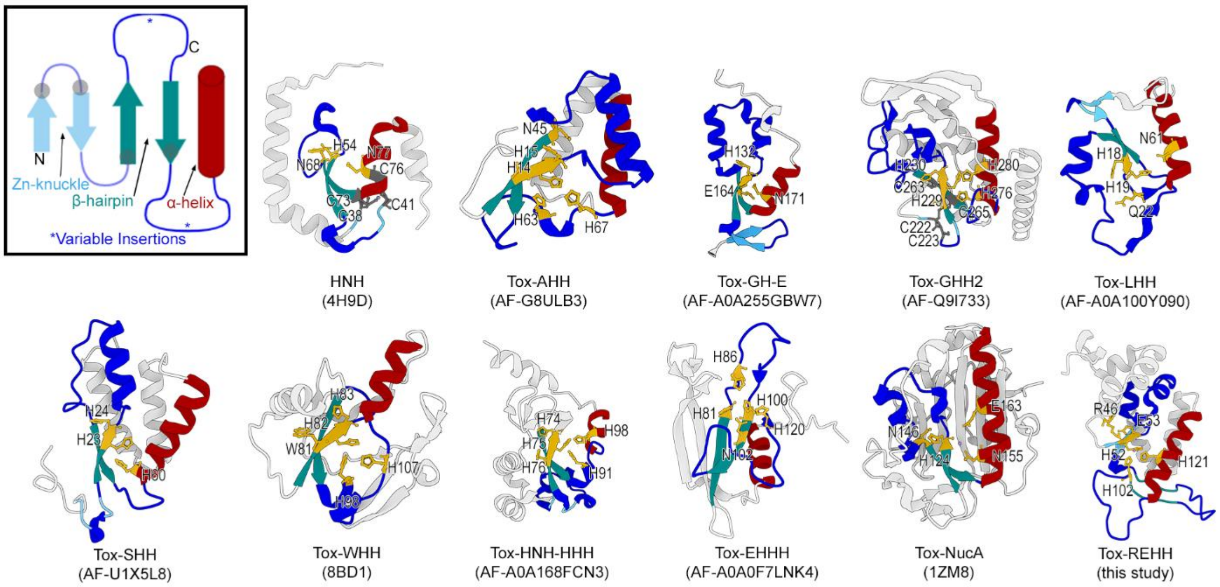


**Figure S3. Rehh preserves the catalytic architecture of the HNH nuclease superfamily.** The topology of the canonical treble-clef motif is shown with its major structural elements highlighted. The β-hairpin (dark cyan) and α-helix (red) are conserved features of HNH nucleases and contain residues that contribute directly to catalysis. The Zn-knuckle (light blue), a defining element of the canonical treble-clef fold, is not universally retained among HNH domains. Likewise, the conserved cysteine residues (gray dots) that typically coordinate $Zn^{2+}$ within the Zn-knuckle are frequently absent in HNH nucleases and can be replaced by alternative metal-coordinating residues, including histidines. Representative HNH toxin domains associated with polymorphic toxin systems are shown with their structural elements colored according to the topology diagram above[1,2]. Conserved catalytic residues are highlighted in yellow, whereas conserved cysteine residues associated with the canonical Zn-knuckle are shown in dark gray when present[3]. The Tox-REHH domain is included for comparison and exhibits the characteristic β-hairpin and α-helix arrangement that defines the catalytic core of the HNH superfamily despite substantial divergence in overall structure. Variable insertions are in dark blue. All protein structures were visualized with Mol*[4].

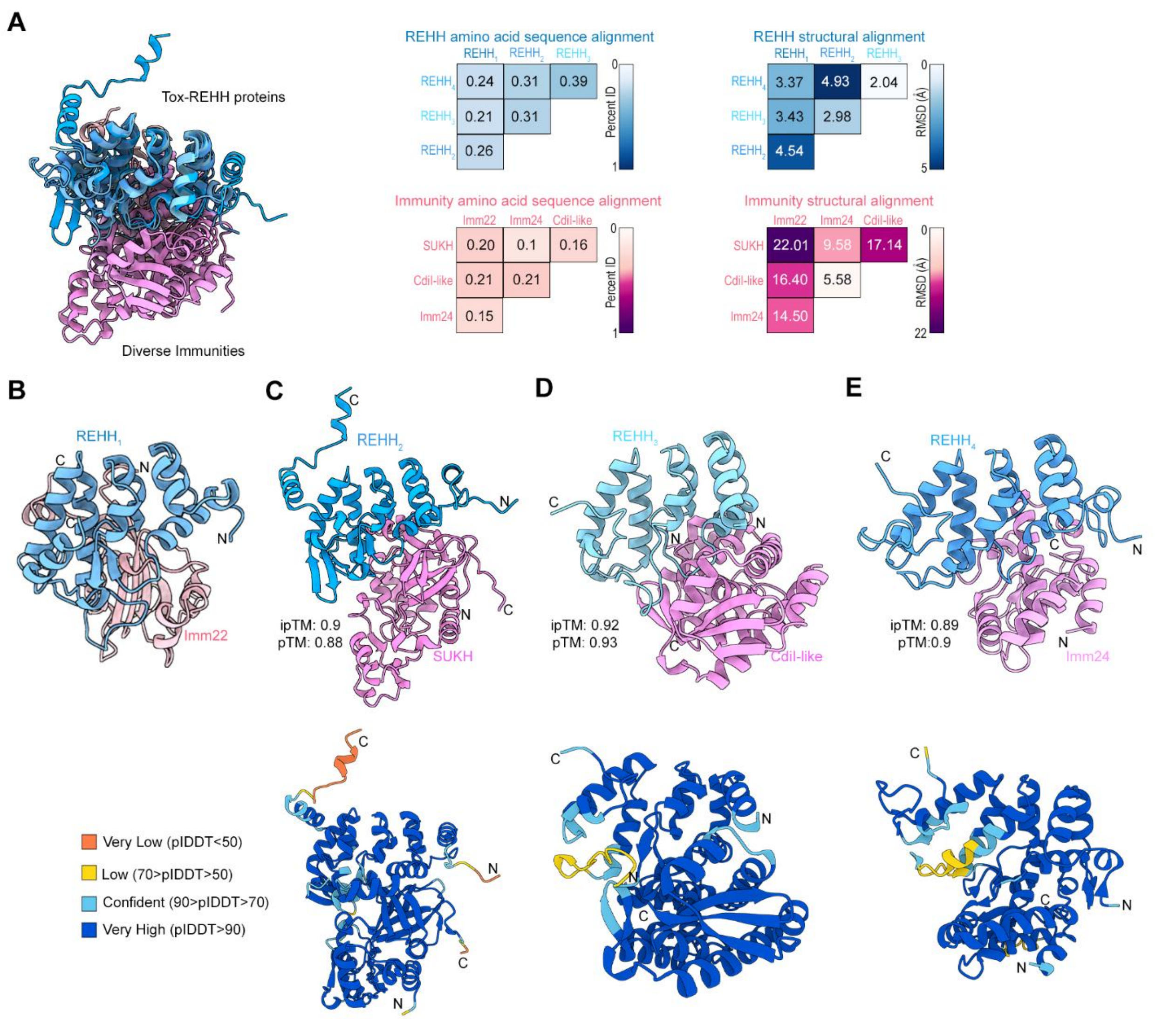


**Figure S4. Amino acid and structural similarities between Tox-REHH homologs and their cognate immunity proteins.** Pairwise sequence and structural comparisons of the Tox-REHH homologs and cognate immunity proteins analyzed in Figure 4. Tox-REHH homologs exhibit limited amino acid sequence conservation but retain a highly similar overall fold. In contrast, the associated immunity proteins display little amino acid or structural similarity and belong to distinct immunity families (Imm22, SUKH, CdiI-like, and Imm24). These comparisons demonstrate conservation of the toxin structural scaffold despite divergence in sequence and immunity-partner architecture. Structure-based alignments were performed using Mol* and figures were generated with ChimeraX[4–7]. Related to Figure 4.

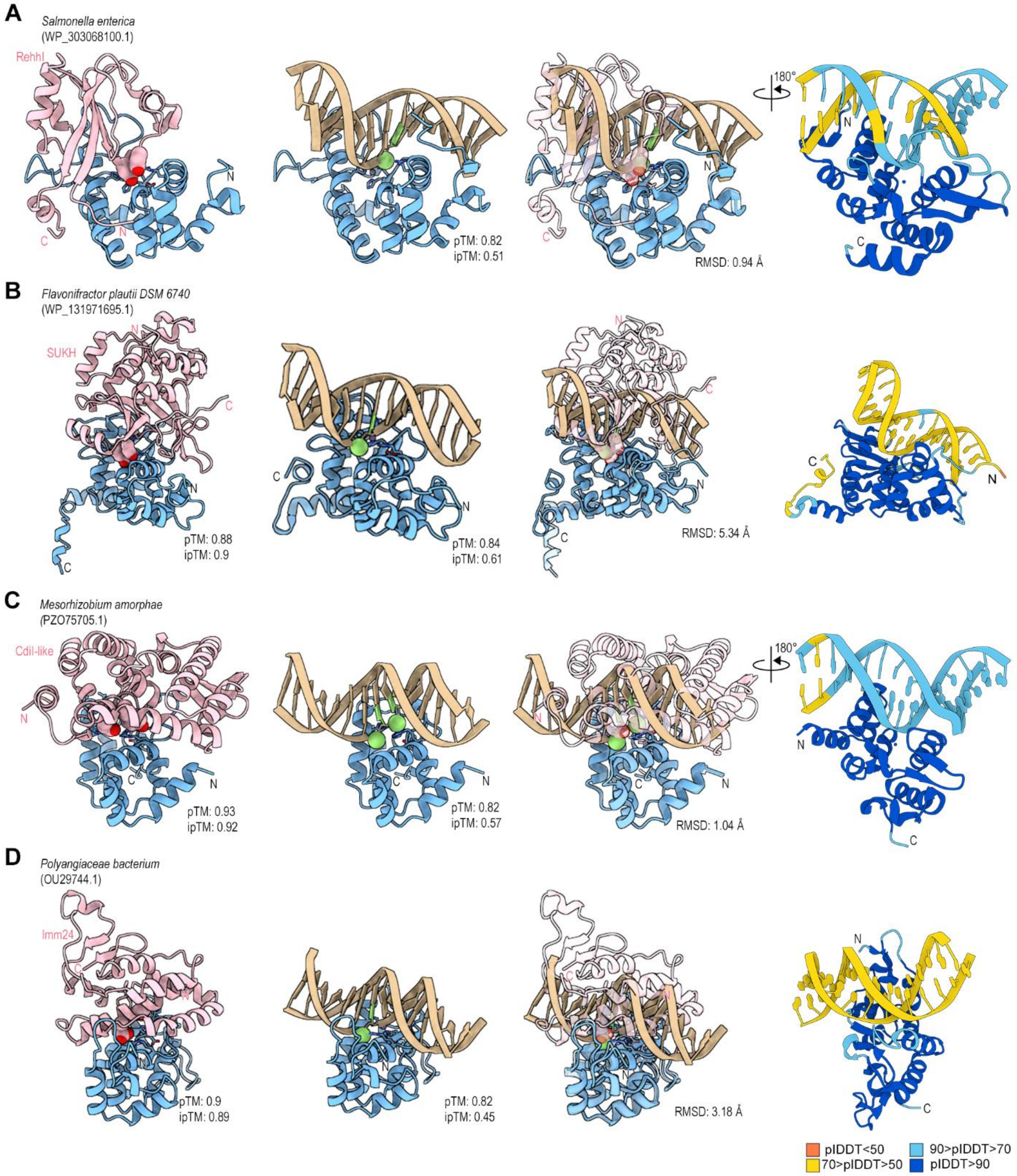


**Figure S5. Conserved acidic residues in immunity proteins occupy positions overlapping predicted DNA-binding sites.** Structural comparison of predicted Tox-REHH-DNA complexes and their corresponding toxin-immunity complexes. The experimentally determined Rehh-RehhI complex is shown as a reference (A). Corresponding analyses are shown for Tox-REHH homologs associated with SUKH (B), CdiI-like (C), and Imm24 (D) immunity families. DNA was modeled into each toxin active site using AF3, and the resulting structures were overlaid with the corresponding toxin-immunity complexes. In all cases, conserved acidic residues from the immunity proteins occupy positions that overlap with phosphates of the modeled DNA backbone, consistent with targeting of the predicted DNA-binding region of the toxin. Related to Figure 4.

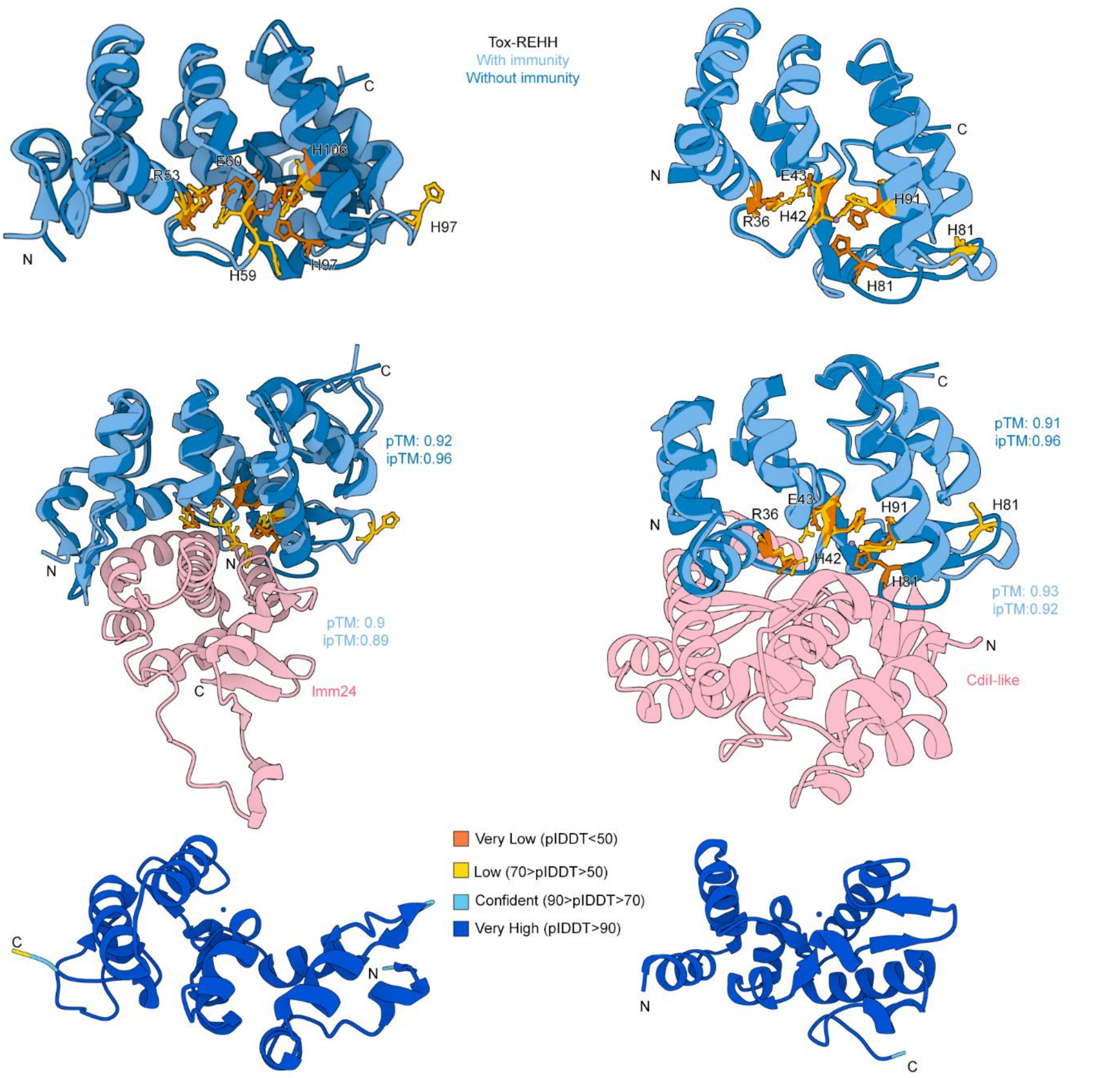


**Figure S6. Structural comparison of Tox-REHH homologs in the presence and absence of cognate immunity proteins.** AF3 models of Tox-REHH homologs predicted in the absence (dark blue) or presence (light blue) of their cognate immunity proteins (pink). Conserved active-site residues are shown in orange (toxin alone) or yellow (toxin-immunity complex). In these cases, in addition to occluding the positively charged active-site region, binding of the immunity protein is associated with a conformational rearrangement of a loop containing a conserved histidine residue. This predicted shift repositions putative catalytic residues relative to the active site and may represent an additional mechanism contributing to toxin inhibition. Structures were visualized using Mol*[4,6]. Related to Figure 4.

**Movie S1. Time-lapse microscopy of *E. coli* carrying plasmid with Rehh under repressing conditions.** Time-lapse microscopy of MG1655 *E. coli* carrying pBRA-Rehh grown under repressing conditions (0.5% D-glucose). Cells maintain normal morphology and growth throughout the experiment. Timestamp is shown as hh:mm. Scale bar, 5 µm.

**Movie S2. Time-lapse microscopy of *E. coli* carrying plasmid with Rehh under inducing conditions.** Time-lapse microscopy of MG1655 *E. coli* carrying pBRA-Rehh grown under inducing conditions (0.2% L-arabinose). Rehh expression results in progressive cell elongation and filamentation over time. Timestamp is shown as hh:mm. Scale bar, 5 µm.

**Table S1. List of species encoding Tox-REHH homologs. Related to Figure 1.**

**Table S2. Gene neighborhoods of Tox-REHH homologs. Related to Figure 1.**

**Table S3. Crystallographic parameters of the Rehh-RehhI complex. Related to Figure 2.**

**Table S4. List of primers used in this study. Extension of the STAR Methods.**
