## Supplementary material for "A widespread bacterial toxin reveals a deeply divergent DNase family neutralized by diverse immunity proteins": Methods

**Key resources table**

| REAGENT or RESOURCE | SOURCE | IDENTIFIER |
| --- | --- | --- |
| Bacterial and virus strains | | |
| *Escherichia coli* DH5α | Lab stock | N/A |
| *Salmonella enterica* serovar Sekondi | Jay Hinton | FD01845383 |
| *Salmonella enterica* serovar Sekondi Δ*tssL* | This paper | N/A |
| *Salmonella enterica* serovar Sekondi Δ*rehh/rehhI* | This paper | N/A |
| *Escherichia coli* MG1655 | Lab stock | N/A |
| *Escherichia coli* SMR 1435 | Shee et al. | N/A |
| *Escherichia coli* BL21 (DE3) | Lab stock | N/A |
| *Escherichia coli* Lemo21 (DE3) | Lab stock | N/A |
| Chemicals, peptides, and recombinant proteins | | |
| Kanamycin | Gibco | cat# 11815-032 |
| Streptomycin | Gibco | cat# 11860-038 |
| Chloramphenicol | Millipore Sigma | cat# C0378 |
| Tetracycline | Millipore Sigma | cat# T7660 |
| Ampicillin | Millipore Sigma | cat# A9518 |
| Bile salts | Oxoid | cat# LP0055J |
| L-rhamnose | Melford | cat# R20600 |
| IPTG (isopropyl-beta-D-thiogalactopyranoside) | Thermo Scientific | cat# R0393 |
| Morpheus | Molecular Dimensions | cat# MD1-46 |
| MemGold2 | Molecular Dimensions | cat# MD1-63 |
| MemGold | Molecular Dimensions | cat# MD1-39 |
| JCSG-plus | Molecular Dimensions | cat# MD1-37 |
| D-glucose | Millipore Sigma | cat# G7021 |
| L-arabinose | Millipore Sigma | cat# 1.01492.0100 |
| SYBR Safe | Invitrogen | cat# S33102 |
| Paraformaldehyde | Millipore Sigma | cat# P6148 |
| DAPI (4',6-diamidino-2-phenylindole) | Thermo Scientific | cat# D1306 |
| Fetal Bovine Serum | Gibco | cat# A5670801 |
| Lysozyme | Thermo Scientific | cat# 89833 |
| Critical commercial assays | | |
| GeneJet plasmid Miniprep kit | Thermo Scientific | K0503 |
| Deposited data | | |
| Rehh-RehhI complex | This paper | PDB:XXXX |
| Oligonucleotides | | |
| See Table S3 | N/A | N/A |
| Recombinant DNA | | |
| pTWIST Rehh | TWIST Bioscience | N/A |
| pTWIST RehhI | TWIST Bioscience | N/A |
| pBRA | Souza et al. | N/A |
| pBRA Rehh | This paper | N/A |
| pBRA RehhR46A | This paper | N/A |
| pBRA RehhE53A | This paper | N/A |
| pBRA RehhH102A | This paper | N/A |
| pBRA RehhH121A | This paper | N/A |
| pEXT22 | Dyxhoorn et al. | N/A |
| pEXT22 RehhI | This paper | N/A |
| pEXT22 RehhID45R | This paper | N/A |
| pEXT22 RehhID47R | This paper | N/A |
| pEXT22 RehhIF69R | This paper | N/A |
| pKD46 | Datsenko & Wanner | N/A |
| pKD3 | Datsenko & Wanner | N/A |
| pCP20 | Datsenko & Wanner | N/A |
| pFPV25.1 | Valdivia & Falkow | N/A |
| pFPV25.1 tssL | This paper | N/A |
| pRSFDuet | Novagen | cat #71341-3 |
| pRSFDuet Rehh RehhI-strep | This paper | N/A |
| pRSFDuet RehhH121A RehhI-strep | This paper | N/A |
| pRSFDuet Rehh-strep RehhI | This paper | N/A |
| pRSFDuet RehhH121A-strep RehhI | This paper | N/A |
| pSC101-P*recA*::GFP | Ronen et al. | N/A |
| Software and algorithms | | |
| jackHMMER | Finn et al. | [HMMER](https://www.ebi.ac.uk/Tools/hmmer/search/jackhmmer) |
| Jalview | Waterhouse et al. | [Jalview Home Page - Jalview](https://www.jalview.org/) |
| MAFFT | Katoh & Standley | [MAFFT - a multiple sequence alignment program](https://mafft.cbrc.jp/alignment/software/) |
| Annotree | Mendler et al. | [AnnoTree](http://annotree.uwaterloo.ca/annotree/) |
| HMMsearch | Eddy | [HMMER](https://www.ebi.ac.uk/Tools/hmmer/search/hmmsearch) |
| ROTIFER | Robson F. de Souza | [GitHub - leepusp/rotifer · GitHub](https://github.com/leepusp/rotifer) |
| GraphPad Prism 10.6.1 | GraphPad Software | [Home - GraphPad](https://www.graphpad.com/) |
| FIJI | Schindelin et al. | [Fiji: ImageJ, with "Batteries Included"](https://fiji.sc/) |
| ChimeraX | Pettersen et al. | [UCSF ChimeraX Home Page](https://www.cgl.ucsf.edu/chimerax/) |
| AlphaFold3 | Abramson et al. | [AlphaFold Server](https://alphafoldserver.com/) |
| Mol* | Sehnal et al. | [Mol* Viewer](https://molstar.org/viewer/) |
| Phenix | Liebschner et al. | [Phenix](https://phenix-online.org/download) |
| CCP4 | Winn et al. | [CCP4 Download](https://www.ccp4.ac.uk/download/index.php#os=windows) |
| COOT | Emsley et al. | [www2.mrc-lmb.cam.ac.uk/personal/pemsley/coot/](https://www2.mrc-lmb.cam.ac.uk/personal/pemsley/coot/) |
| REFMAC5 | Murshudov et al. | [www2.mrc-lmb.cam.ac.uk/groups/murshudov/content/refmac/refmac.html](https://www2.mrc-lmb.cam.ac.uk/groups/murshudov/content/refmac/refmac.html) |
| Other | | |
| Nikon HCA Microscope | Microscopy & Flow Cytometry Core Facility (UT Austin) | N/A |
| Nitrocellulose Membranes (0.2 µm) | Bio-Rad | cat# 1620112 |
| HisTrap HP 5 mL column | Cytiva | cat# 17524802 |
| Hiload 26/600 superdex 75 pg gel filtration column | Cytiva | cat# 28989334 |
| Mosquito liquid handling system | SPT LabTech | N/A |
| Agilent BioTek Synergy H1 Plate Reader | Agilent | N/A |
| ChemiDoc Imaging System | Bio-Rad | cat# 12003153 |
| Streptrap XT 1 mL column | Cytiva | cat# 29401320 |
| Histrap HP 1 mL column | Cytiva | cat# 17524701 |
| Amicon ULTRA-15 centrifugal filter (10 kDa) | Millipore Sigma | cat# UFC901024 |
